# *Russula* (Basidiomycota, Russulales, Russulaceae) subsect. *Roseinae* “down under”

**DOI:** 10.1101/2023.08.18.553850

**Authors:** Bart Buyck, Egon Horak, Jerry A. Cooper, Yu Song

## Abstract

The present contribution presents species of *Russula* subsect. *Roseinae* Sarnari from the southern hemisphere. *Russula incrustata* Buyck, sp. nov. and *R. koniamboensis* Buyck, sp. nov. are described from New Caledonia, *R. purpureotincta* R. F. R. McNabb from New Zealand is redescribed in detail and two secotioid species, *R. albobrunnea* T. Lebel from Australia and *R. kermesina* (R. F. R. McNabb) T. Lebel from New Zealand are shown to be the first known secotioid taxa in *Roseinae*. The systematic placement and importance of these southern taxa is discussed.

## Introduction

Extant species of *Russula* subsect. *Roseinae* have been suggested to have diverged about three to one million years ago (Looney et al., 2020) and are part of the most diverse, major lineage of the genus in the northern hemisphere, viz. the Crown clade of subgenus *Russula* (Buyck et al. 2018). Species in subsect. *Roseinae* are morphologically characterized by the pink, red or whitish to yellowish pileus, the white to pale cream spore print, the predominantly mild taste, the context and lamellae turning eosin red with sulfovanillin and the lack of typical gloeocystidia in pileipellis and context. In addition, the involved species possess “primordial hyphae”, i.e. hyphal terminations with acid-resistant incrustations (see Romagnesi 1967: 58-59) which are generally emerging above the other hyphal extremities. All *Roseinae* also have inflated cells in the lower pileipellis composing a more or less well-developed pseudoparenchymatic lyer from which the hyphal terminations originate.

In Europe, *Russula* subsect. *Roseinae* is represented by merely two widely accepted species, *R. velutipes* Velen. (syn.: *R. rosea* Quél.) and *R. minutula* Velen., but the subsection is much more diverse in other regions of the northern hemisphere. Indeed, recent taxonomic studies on *Roseinae* report seven described North American species and an additional six species that are still not formally described (Looney et al., 2020; Manz et al., 2021): *R. cordata* Looney*, R. rheubarbarina* Looney*, R. rubellipes* Fatto*, R. peckii* Singer*, R. pseudopeckii* Fatto and one undescribed relative, *R. cardinalis* Looney*, R. albida* Peck (syn.: *R. purpureomaculata* Shaffer) with two undescribed American relatives, one undescribed *R. minutula* aff, as well as two undescribed species in the ‘*magnarosea*’-lineage). In addition, four more species of *Roseinae* were described from the Central American montane forests in Western Panama (Manz et al., 2021: *R. cornicolor* Manz & F. Hampe*, R. zephyrovelutipes* Manz & F. Hampe*, R. oreomunneae* Manz, F. Hampe & Corrales and *R. cynorhodon* Manz & F. Hampe), now bringing the total of putative American species in *Roseinae* (s. l.) to 17 taxa. These same authors (Manz et al. 2021) confirmed or suggested the position of the following thirteen Asian phylogenetic species in *Roseinae*: *R. dhakuriana* K. Das, J. R. Sharma & S. L. Mill.*, R. hakkae* G. J. Li, H. A. Wen & R. L. Zhao*, R. kewzingensis* K. Das, D. Chakr. & Buyck with one undescribed relative, *R. guangxiensis* G. J. Li, H. A. Wen & R. L. Zhao with one undescribed sister species, two undescribed Asian relatives to *R. velutipes*, two more undescribed Asian species close to *R. pseudopeckii*, two undescribed Asian relatives to *R. peckii*, as well as one sister species to *R. cynorhodon*.

*Russula rimosa* Murrill, again a North American species, is only known from the type collection and was classified in subsect. *Roseinae* based on its morphology (Adamčík & Buyck, 2012), but so far, any attempt of sequencing the DNA of this species has been unsuccessful. Sequence data are also needed for the Indian *R. sharmae* K. Das, Atri & Buyck, another potential member of subsect. *Roseinae* based on its microscopic features, but it is unusual in producing an almost yellowish spore print (Das et al., 2013). This brings the total of potential northern hemisphere *Roseinae* to 30 recognized species or about five times more than hardly a few years ago !

Looney et al. (2020) suggested a northern hemisphere distribution for *Roseinae* and inferred an Appalachian origin for the subsection in North America, followed by in situ diversification of these species in the Appalachian Mountains roughly since the mid-Miocene. Whereas published sequence data had already suggested the existence of several potential *Roseinae* in Oceania or Australasia (Buyck et al., 2018; Lebel & Tonkin, 2007; Cooper & Leonard, 2014: Cooper, 2021), none of these southern species has ever been discussed in the literature. The aim of this study is to identify the actual taxonomic status and phylogenetic position of these southern hemisphere relatives.

## Materials and Methods

### Morphological study

Fresh fruiting bodies were photographed in the field and in the lab, descriptive notes and spore prints were taken and tissue samples transferred in CTAB solution for subsequent sequencing.

Micromorphological characters were studied using a Nikon Eclipse E400 microscope under oil-immersion lens at a magnification of x1000. All drawings of microscopical structures were made with a ‘camera lucida’ using a Nikon Y-IDT drawing attachment at a projection scale of x2400. Contents of hymenial cystidia and pileocystidia in the figures are indicated schematically, with the exception of a few elements where contents are indicated as observed in Congo red preparations from dried material. Spores were observed in Melzer’s reagent. All other microscopic observations were made in ammoniacal Congo red, after a short aqueous KOH pre-treatment near boiling temperature to improve tissue dissociation through gelatinous matrix dissolution. All tissues were also examined for the presence of ortho-or metachromatic contents or incrustations in Cresyl blue as explained in Buyck (1989).

### DNA extraction, PCR and sequencing

Fungal genomic DNA was isolated from fresh material stored in cetyl-trimethyl-ammonium bromide buffer (CTAB 1x). Five loci were tentatively amplified: 900–1400 base pairs of the ribosomal nuclear large subunit (nucLSU) using primers LROR and LR7; 600 base pairs of the ribosomal mitochondrial small subunit (mitSSU) with primers MS1 and MS2 (White et al. 1990); 1300 base pairs of the largest subunit of the RNA polymerase II (RPB1) with primers RPB1-AF (Stiller & Hall, 1997) and RPB1-CR (Matheny et al., 2009); 700 base pairs of the second largest subunit of the RNA polymerase II (RPB2) using primers RPB2-6F and fRPB2-7cR (Liu & Hall, 2004) and 900 base pairs of the translation elongation factor 1-alpha (TEF1) using primers EF1-F and EF1-R (Morehouse et al., 2003). Amplifications were performed under the conditions and with the reagents of the Taq PCR core kit (QIAGEN, Inc., Valencia, California, USA). Sequencing used the amplification primers, reagents and conditions of the BigDye Terminator v3.1 Cycle sequencing Kit and an automated capillary sequencer ABI 3700 DNA analyzer (Perkin Elmer, Applied Biosystems, Foster City, CA, USA). Most sequences for these various loci were obtained for *R. incrustata* and have already been deposited in GenBank (as *R. roseinae sp. VH-2016n strain 735/BB 09.172*) and published in the context of a *Russula* world phylogeny (Buyck et al. 2018). The newly published sequences have all been deposited in GenBank (www.ncbi.nlm.nih.gov), viz. ITS (OM397456-OM397459) and tef1 (OM365994-OM365996, OM370807) for both new species from New Caledonia, eight ITS sequences for *R. purpureotincta* R.F.R.McNabb (OR348209, OR348217, OR348277-OR348281), including one from the holotype, and a single ITS sequence for *R. kermesina* (R.F.R.McNabb) T.Lebel (OR348284).

### Phylogenetic analysis

Separate phylogenetic analyses based on ITS and *tef1* performed with Maximum Likelihood method were chosen in function of available sequence data for the southern species. The sampling of northern hemisphere taxa is based on Manz et al. (2021). Each dataset was automatically aligned by MAFFT v 7.427 (Katoh & Standley, 2013), then manually adjusted and trimmed with BioEdit v7.0.9 (Hall, 1999). The final ITS alignment consisted of 70 sequences and comprised 750 bp ; the *tef1* alignment was 921 bp long (excluding introns) and comprised 41 sequences, *Russula emeticicolor* (Jul.Schäff.) Singer and *R. lilacea* Quél. belonging to subsect *Lilaceinae* (Melzer & Zvára) Jul.Schäff. were chosen as the outgroup. A rapid bootstrapping (BS) algorithm of 1000 replicates was executed in RAxML 7.2.6 (Stamatakis, 2006), followed by a heuristic ML search for the best tree using the GTRGAMMA model. All parameters in RAxML analysis were kept at default. Bootstrap value (BS) exceeding or equal to 70% was considered to represent significant support.

## Results

### Phylogeny

The ITS phylogeny (Fig. 1) comprises 68 sequences representative of taxa in *Roseinae* and two sequences of *Lilaceinae* chosen as out-group. Southern *Roseinae* are represented by six previously deposited sequences in GenBank and twelve newly generated sequences (see above). In this phylogeny, the three New Caledonian specimens of *R. koniamboensis* are placed sister with significant, although moderate support (MLbs=76%) to our second new species from New Caledonia, *R. incrustata*. Both New Caledonian species are again placed sister with full support (MLbs=100%) to a clade composed of all sequenes of *R. purpureotincta* from New Zealand (one was previously deposited under the wrong name of *R. cremeoochracea* in GenBank). It is for the very first time that we also place two secotioid species in subsect. *Roseinae*: *R. kermesina* from New Zealand and *R. albobrunnea* from Australia. For both, only ITS sequence data are available and our analysis reveals that these two secotioid taxa are monophyletic with full support (MLbs=100%). We obtained no significant support to place this clade in relation to any other particular clade, but our phylogeny suggests close relationship with a clade composed of *R. pseudopeckii*, *R. zephyrovelutipes* and related taxa.

**Fig. 1.**
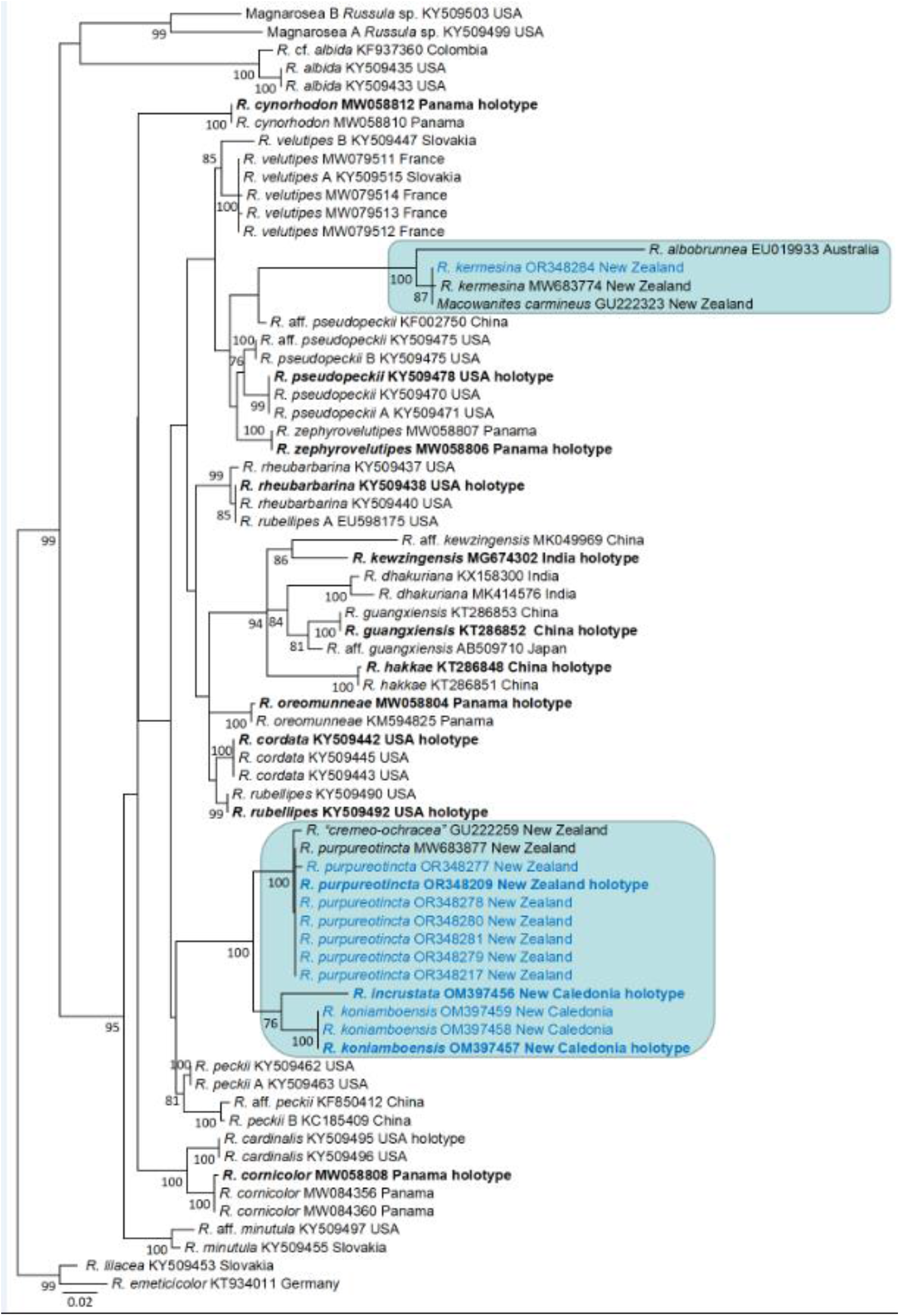
ITS phylogeny of *Russula* subsect. *Roseinae*. Southern hemisphere species are indicated by coloured rectangles. Type specimens are indicated in bold.

The *tef1* phylogeny (Fig. 2) comprises 41 sequences and places only the New Caledonian species as no *tef1* sequences are available for the other southern species. Whereas the exact relationships of both New Caledonian species with the other species in the subsection remained unresolved in the ITS phylogeny (fig. 1), the *tef1* phylogeny offers now strong support (MLbs=97%) to put both New Caledonian sister to the /cardinalis lineage as delimited in Manz et al. (2021). The latter lineage comprises merely two species, viz. *R. cornicolor* associated with *Oreomunnea* Oerst. (fam. Juglandaceae) in Western Panama and *R. cardinalis* associated with *Quercus* L. in the Appalachian Smoky Mts of Tennessee, USA.

**Fig. 2.**
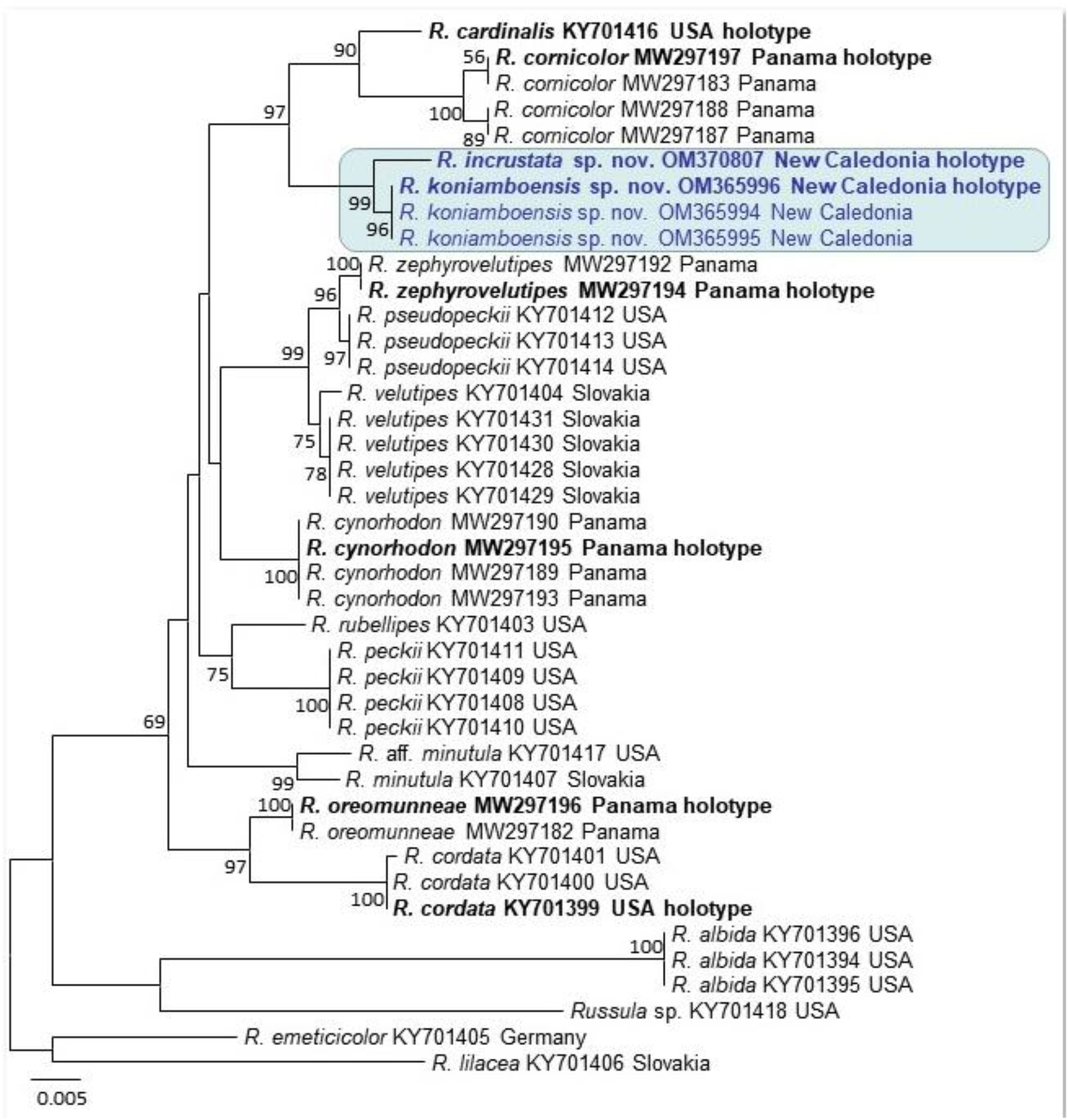
The *tef1* phylogeny of *Russula* subsect. *Roseinae* placing both New Caledonian species (blue rectangle). Type specimens are indicated in bold.

**Fig. 3.**
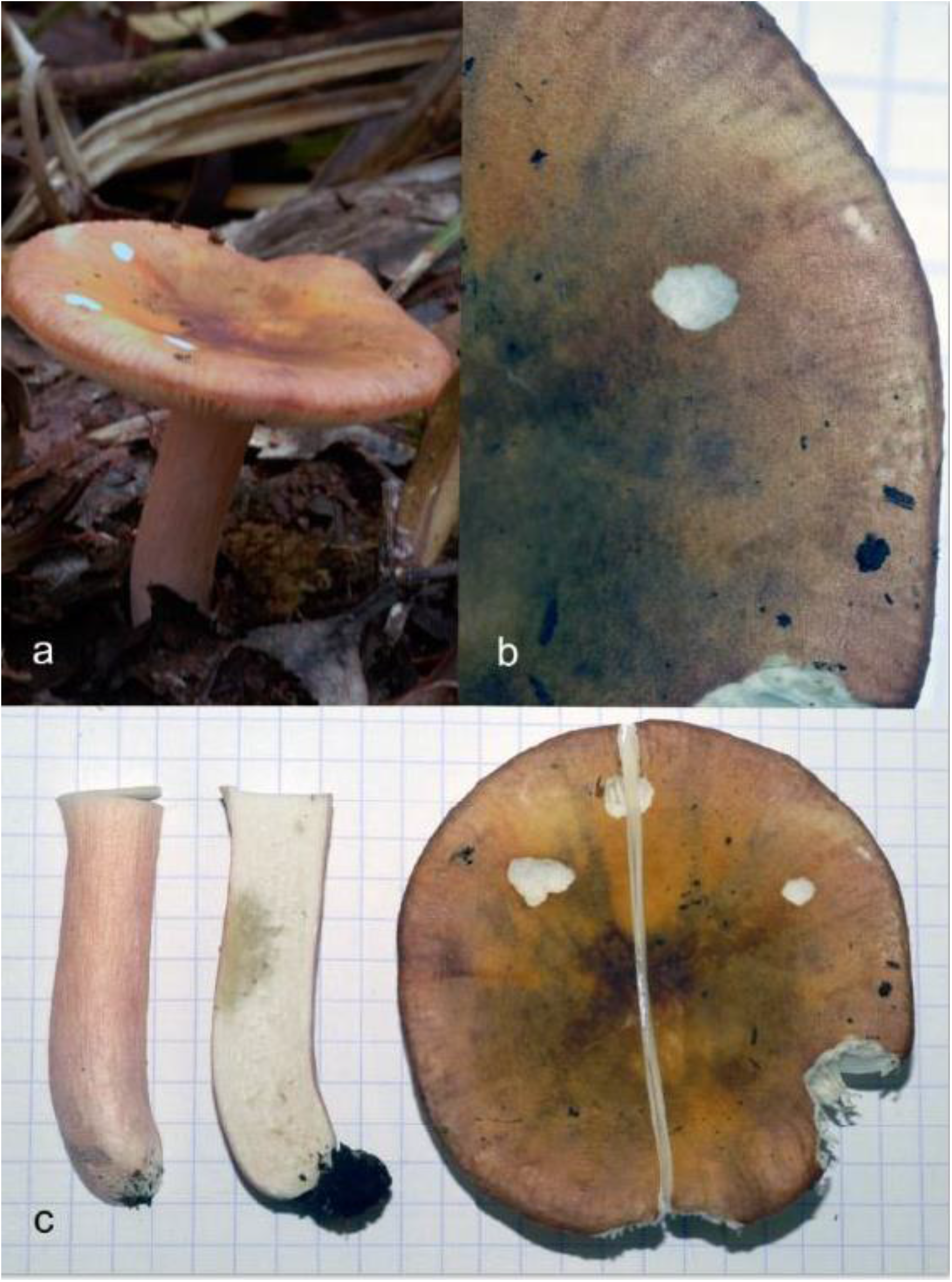
*Russula incrustata* Buyck, sp. nov. (holotype): Fresh basidiomata. a, in situ; b, detail of the pileus surface; c, view of pileus and stipe surface. Photo credit: B. Buyck.

Considering the topologies obtained in our ITS and *tef1* analyses and those obtained in the abovementioned papers with multigene analyses (see introduction), it seems likely that southern hemisphere *Roseinae* occur in at least two different lineages within the subsection: the /cardinalis-lineage for the agaricoid species from New Caledonia and New Zealand, and a mixed American-Asian lineage for both secotioid taxa.

#### Taxonomy

In the below paragraphs we will provide descriptions for the two new New Caledonian species, as well as a description for the here newly reported collections for the related *R. purpureotincta* from New Zealand.

### 1. Russula incrustata Buyck, sp. nov

Figs.: 3, 5-8, 9d

Differs from *R. koniamboensis* Buyck, sp. nov. in the vividly colored pileus, pinkish stipe, the somewhat smaller spore size and its occurrence under endemic *Arillastrum gummiferum*.

Holotype. — NEW CALEDONIA: Les Bois du Sud, near blockhut, S22.171713-E166.760103, ca. 300 m alt., on ultramafic soil in monodominant rain forest of endemic *Arillastrum gummiferum* (Brongn. & Gris) Pancher ex Baill. [fam. Myrtaceae], 27 March 2009, *735/BB 09.172* (PC0142414).

Mycobank. — xxxx

Genbank. — OM370807 (ITS), KU237588 (LSU), KU237436 (mitSSU), KU237873 (*rpb2*), KU238015 (*tef1*), KU237728 (*rpb1*)

Etymology. — the epithet refers to the distinct incrustations present on the cell walls of the lower pseudoparenchyma of the pileipellis and other parts of the context.

### Pileus

Medium-sized, 64 mm diam., plane or very gently and widely depressed in the center, inconspicuously striate near margin; surface peeling 2/3 radius, dull, not hygrophanous, felty-velutinous, never viscid, not fragmenting in areolae or squamae but slightly concentrically pruinose, colour range from pale brown, orange, pink, lilac, purple or vinaceous, not paler in the center with age.

### Stipe

51 × 11-12 mm, central, cylindrical; surface smooth, slightly longitudinally wrinkled, not pruinose, entirely pinkish but white at the extreme base, firm, context soft spongy without cavities, basal mycelium absent.

### Lamellae

Equal in length, adnate, without anastomoses or forks, brittle, 8 mm high, off-white.

### Context

Brittle, white, colour unchanging on injury or with age, about 6 mm thick in pileus above gill attachment to stipe, with FeSO4 hardly changing color, merely faintly grey inside stipe and weakly pinkish on stipe surface, insensitive to Guaiac reaction negative. Taste and odor not distinctive.

### Spore print

White.

### Spores

(6.46)6.89**–7.27**–7.65(7.92) x (5.42)5.67–**5.92**–6.17(6.46) μm, broadly ellipsoid, Q = (1.10)1.17–**1.23**–1.29(1.36); ornamentation subreticulate, composed of large, prominent, moderately distant, conical and strongly amyloid spines, up to 1.5 μm high, connected by frequent fine lines into an incomplete network; suprahilar spot well–developed, varying from strongly amyloid to verruculose, grayish and poorly amyloid.

### Basidia

30–40(–58) × 11–15 μm, narrowly to broadly clavate, 4–spored with stout sterigmata; basidiola clavate.

*Subhymenium*

*pseudoparenchymatic.*

### Hymenial gloeocystidia

On lamella sides mostly 50–74 × 8–10 μm, clavate to fusiform, frequently mucronate to appendiculate at apex, up to 10 μm long, rarely obtuse rounded, originating in subhymenium and longer as basidia, walls up to 2(–2.5) µm thick; contents mainly restricted to some refringent inclusions at apex, not reacting to sulfovanillin, rarely with up to 5 secondary septa; cystidia near the lamellae edges smaller, up to 30 μm long.

### Marginal cells

Occupying most of the lamellar edges, sitting on 1–2 basal cells, mostly 13–29 × (3–)5–9 µm, very variable in shape, several reminding of the terminal cells in the suprapellis (but smaller), but usually with 1–4 diverticulate pointed to obtuse–rounded outgrowths.

### Pileipellis

Two–layered, not distinctly delimited from the underlying context; subpellis composed of a 40–60 μm thick, loose pseudoparenchyma composed of intertwined and strongly ramifying, ascending to erect, irregularly shaped cells, in the lower part with very distinct orthochromatic zebroid incrustations, larger inflated cells at the base up to 12(–15) µm wide, giving rise to smooth–walled 2–4(–5) narrower cells at the pileus surface; subterminal cells frequently branched, thin–walled, barrel–shaped or subcylindrical; terminal cells mostly longer in comparison, (8–)16–23(–39) × 2–4(6) μm, often undulated in outline, typically attenuating towards a minutely capitate apex or with one subapical lateral diverticulum. Primordial hyphae difficult to distinguish in shape from other extremities, recognizable at the terminal cell measuring 15–31 x 3–5 µm and filled with refringent, granular-heteromorphous contents in their very upper part. Cystidioid hyphae absent from subpellis and context. Oleiferous hyphae or hyphal fragments abundant in subpellis.

### Clamp connections

absent.

*Notes*: *Russula incrustata* is a beautiful and colourful species that we found only once. There are no environmental sequences available for it, which could indicate that this taxon is rather rare. Perhaps its ectomycorrhizal association with the rare and endemic *Arrilastrum gummiferum* is the explanation. In the multigene phylogenetic analysis of worldwide *Russula* (Buyck et al., 2018), this species was already clearly placed in *Roseinae* as “R.roseinae sp. ined”.

Scale bar = 10 µm.

### 2. Russula koniamboensis Buyck, sp. nov

**Figs. 4, 9a-c, 10-14**

**Fig. 4.**
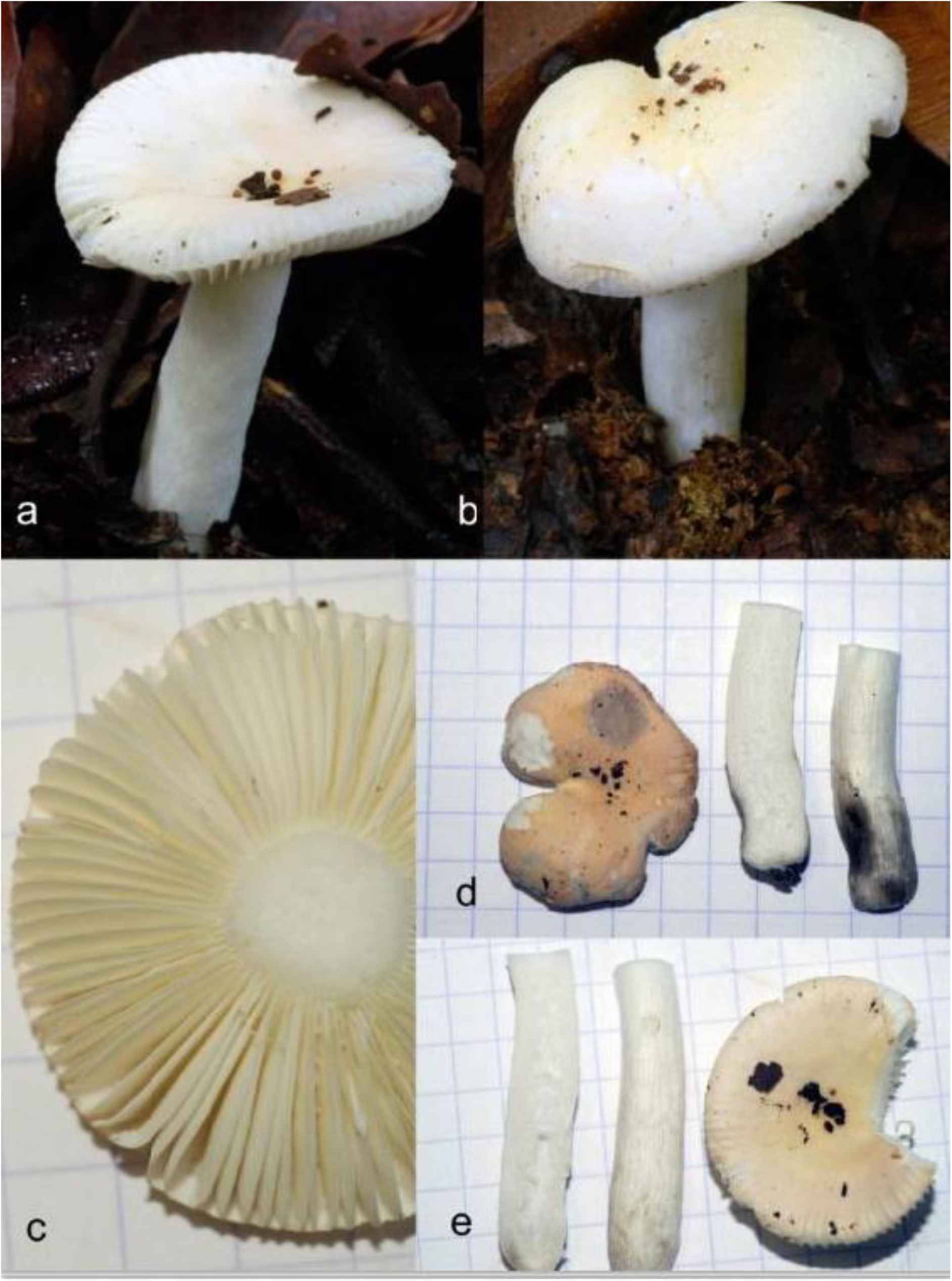
*Russula koniamboensis* Buyck, sp. nov. Fresh basidiomata. a-c, holotype; d-e, 730/BB09.170. Photo credits: B. Buyck.

**Fig. 5.**
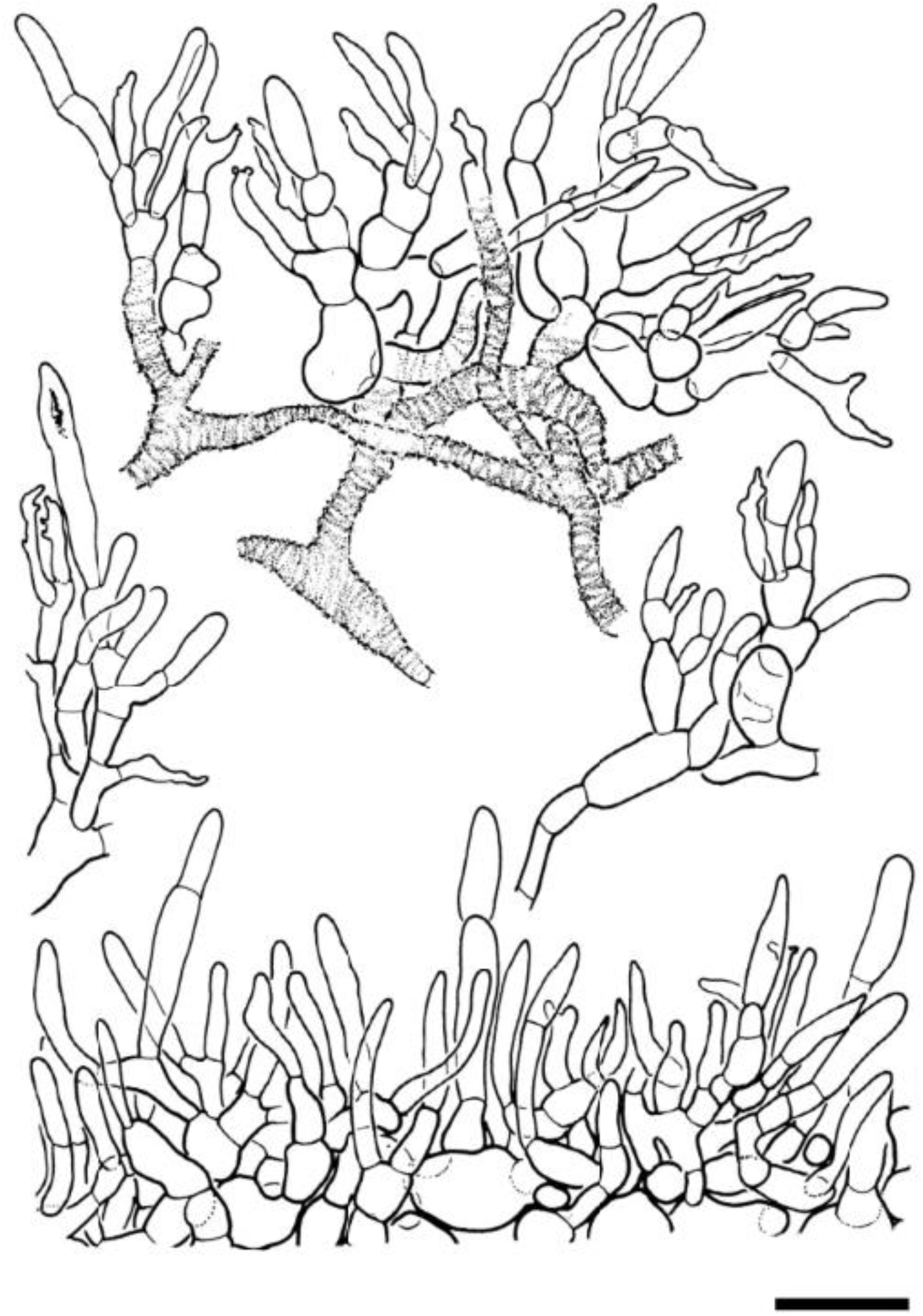
*Russula incrustata* Buyck, sp. nov. (*735/BB09.172*, holotype): Details of the pileipellis. - Scale bar = 20 **µm**. Drawings B. Duhem.

**Fig. 6.**
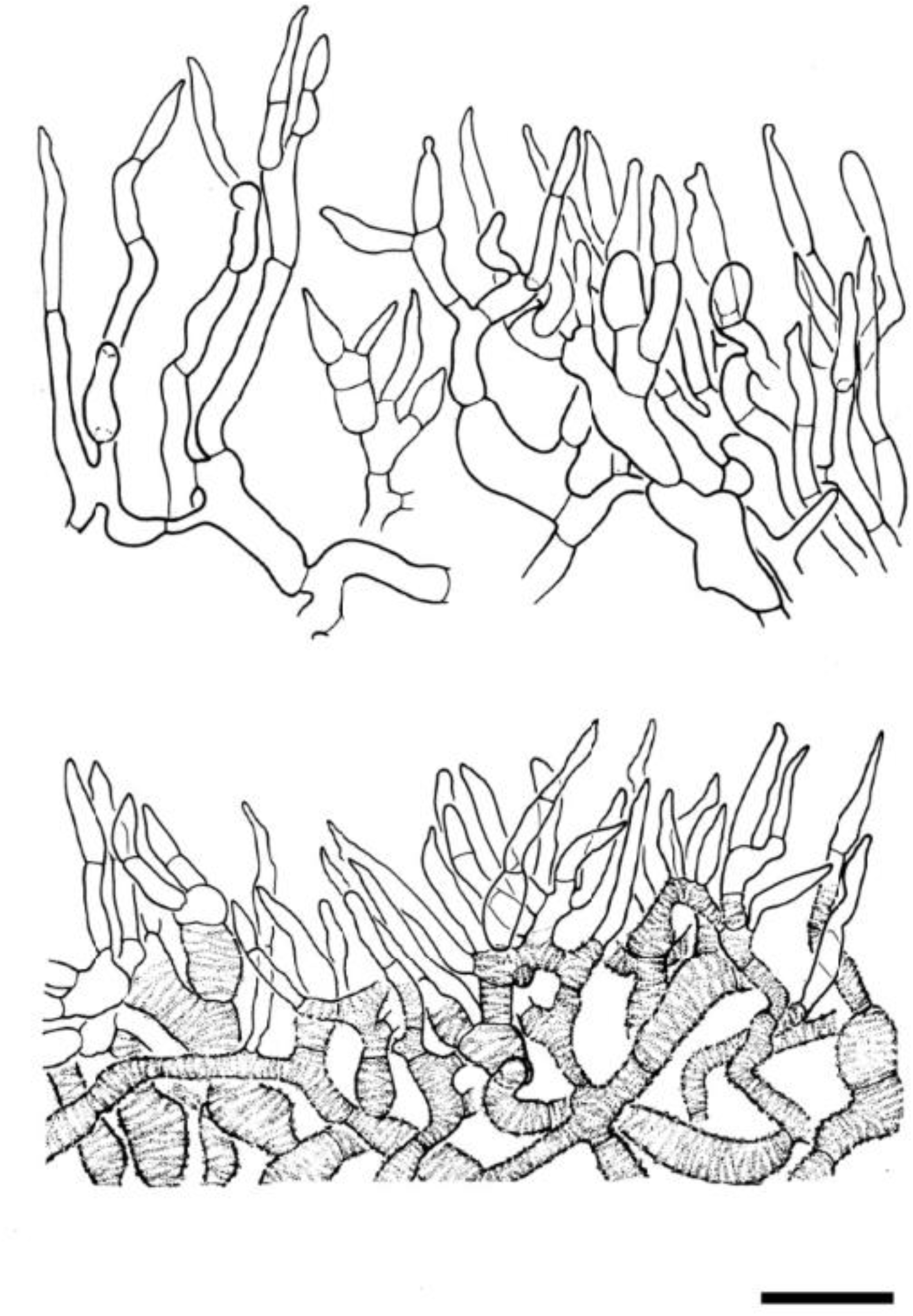
*Russula incrustata* Buyck, sp. nov. (*735/BB09.172*, holotype): Hyphal terminations of the pileipellis, continued. - Scale bar = 20 µm. Drawings B. Duhem.

**Fig. 7.**
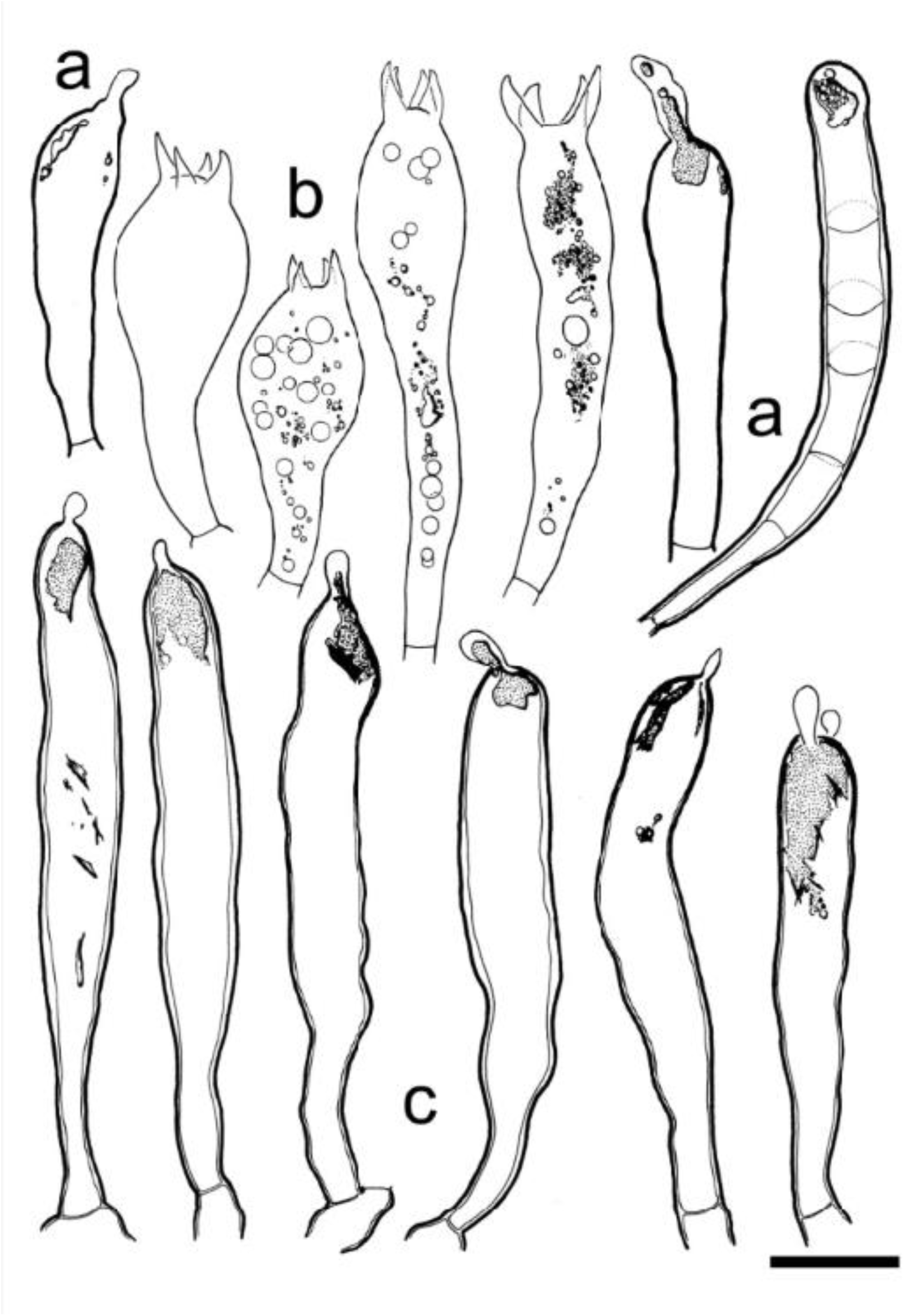
*Russula incrustata* Buyck, sp. nov. (*735/BB09.172*, holotype): Details of the hymenium: a, cheilogloeocystidia, b, basidia; c, gloeocystidium with secondary septa; d, pleurogloeocystidia. - Scale bar = 10 µm. Drawings B. Duhem.

**Fig. 8.**
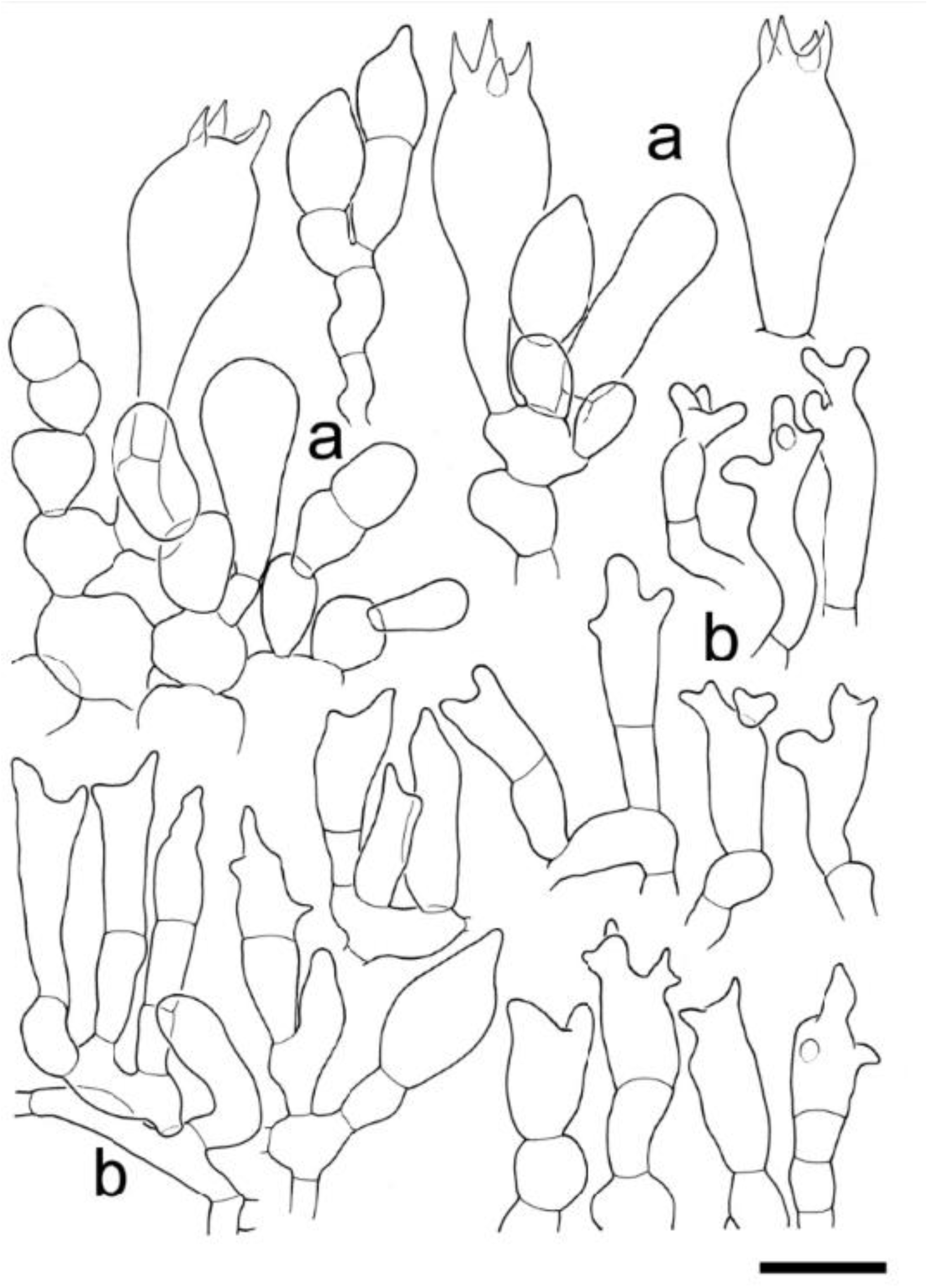
*Russula incrustata* Buyck, sp. nov. (*735/BB09.172*, holotype). Details of the hymenium near the gill edge: a, basidiola and basidia; b, marginal cells. - Scale bar = 10 µm. Drawings B. Duhem.

**Fig. 9.**
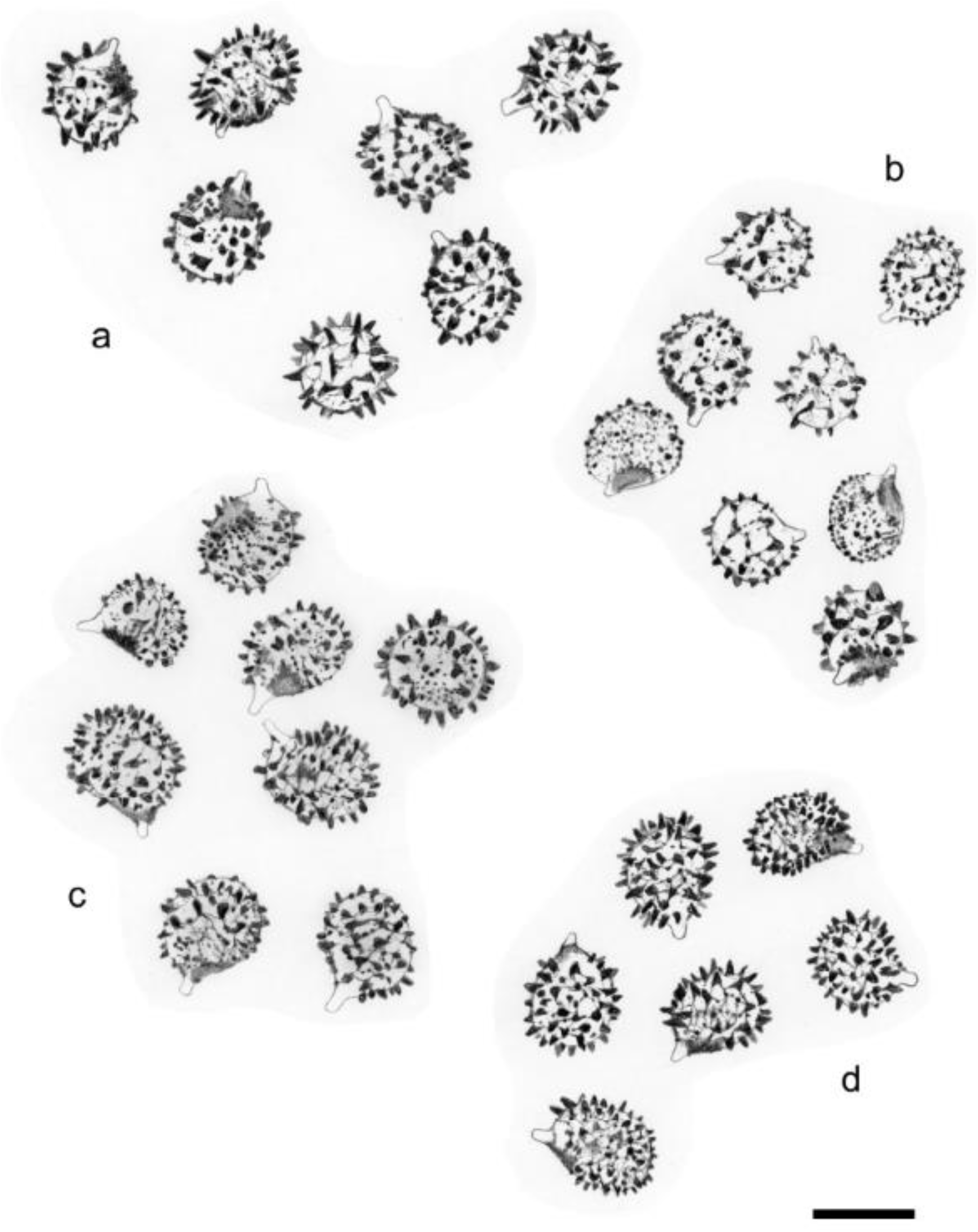
Spore clouds. a-c. *Russula koniamboensis* Buyck, sp. nov. (a: *730/BB09.11,* b: *742/ BB09.346*, c: *722/ BB09.022*, holotype). d, *Russula incrustata* Buyck, sp. nov. (holotype).

**Fig. 10.**
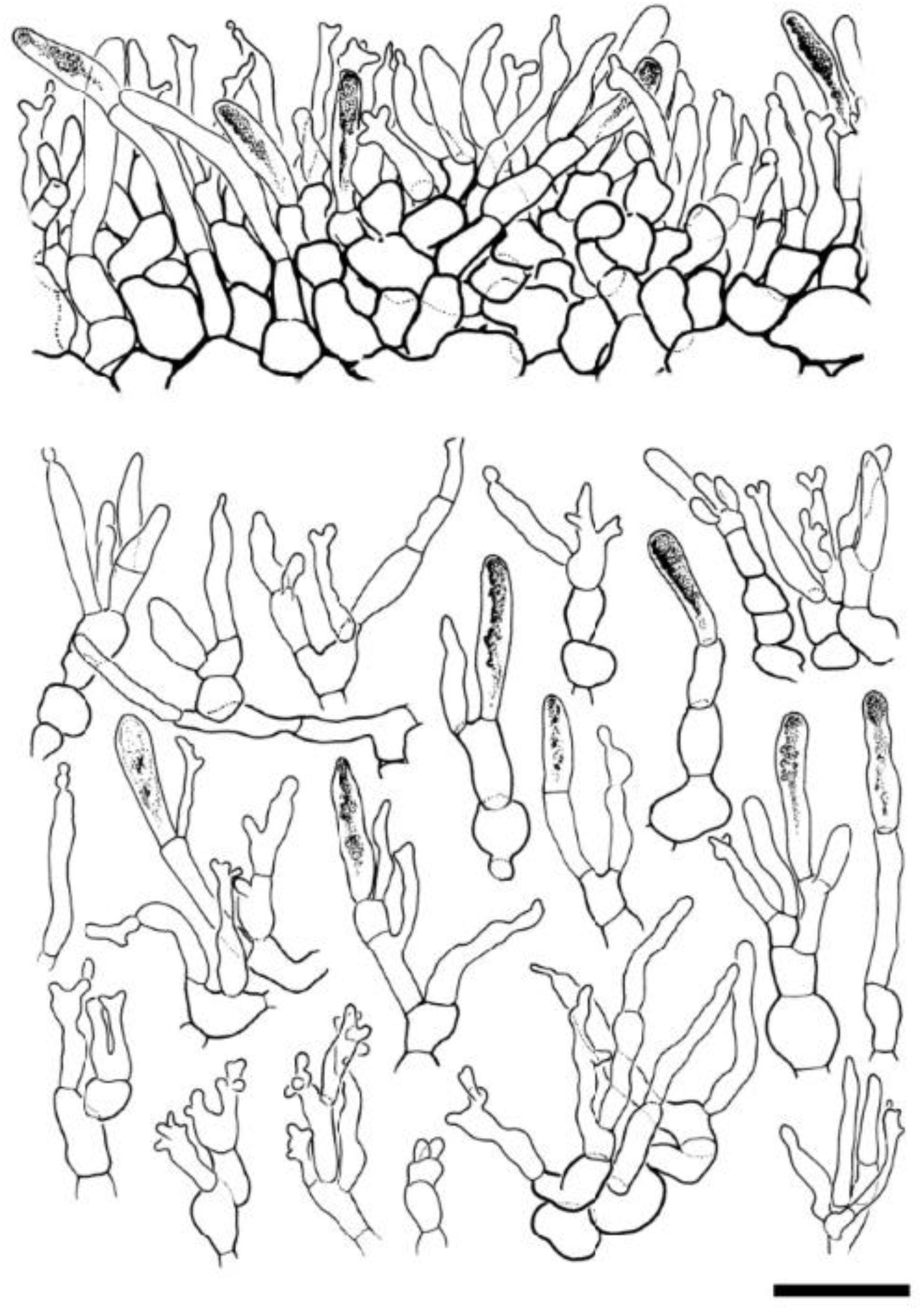
*Russula koniamboensis* Buyck, sp. nov. (*722/BB09.022*, holotype). Details of the pileipellis. Note that all capitate-mucronate endings are optically empty; these are not pileocystidia, notwithstanding they are morphologically very similar to the pileocystidia of species in subg. *Heterophyllidiae*, while all terminal cells of primordial hyphae are obtuse-rounded at their apex and possess refringent contents reminiscent of typical pileocystidia. Scale bar = 20 µm. Drawings B. Duhem.

**Fig. 11.**
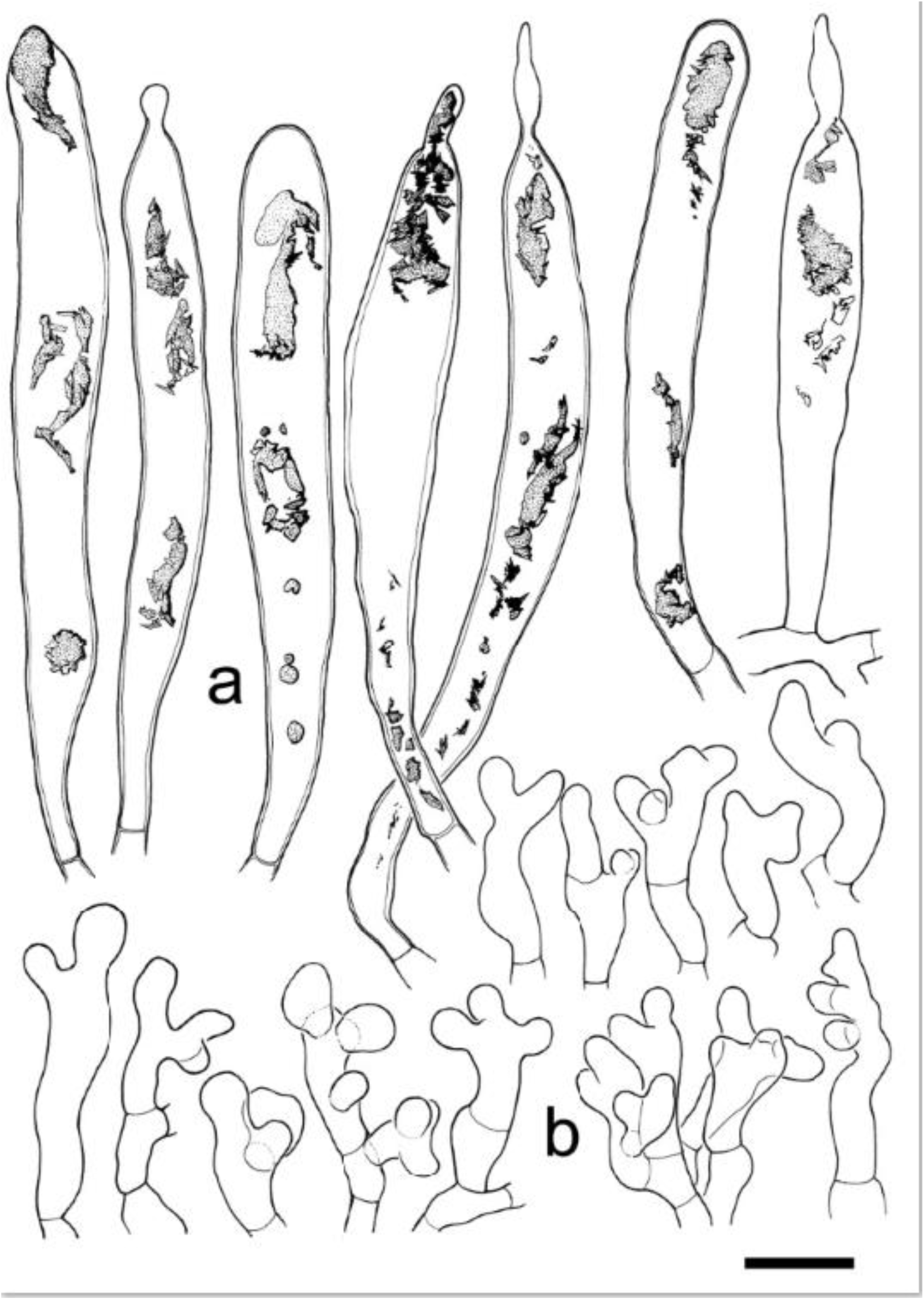
*Russula koniamboensis* Buyck, sp. nov. (*722/BB09.022*, holotype). Elements of the hymenium; top row: pleurogloeocystidia with indication of contents; bottom, marginal cells. Scale bar = 10 µm. Drawings B. Duhem.

**Fig. 12.**
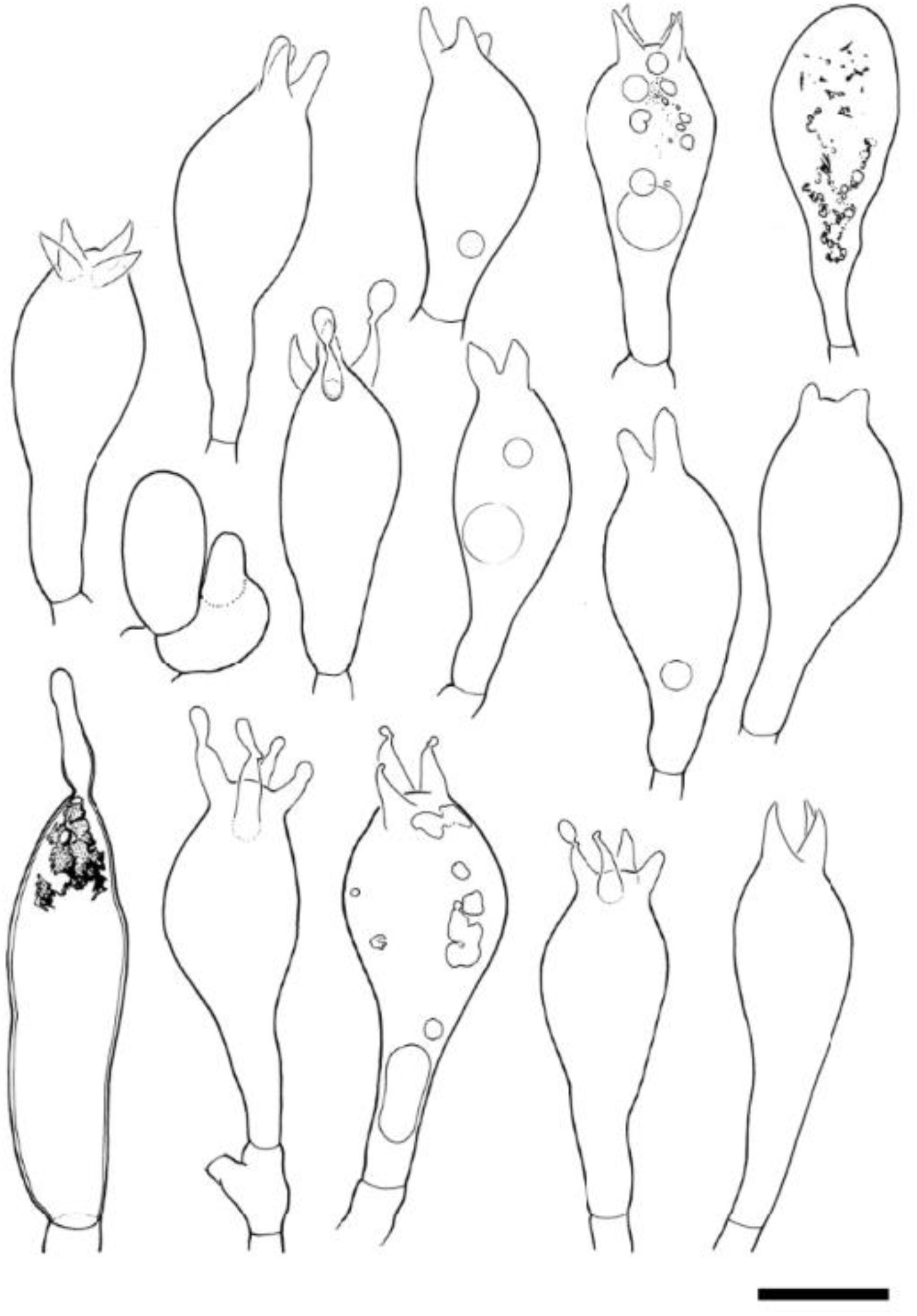
*Russula koniamboensis* Buyck, sp. nov. (*722/BB09.022*, holotype). Elements of the hymenium; cheilogloeocystidium; basidia and basidiola. - Scale bar = 10 µm. Drawings B. Duhem.

**Fig. 13.**
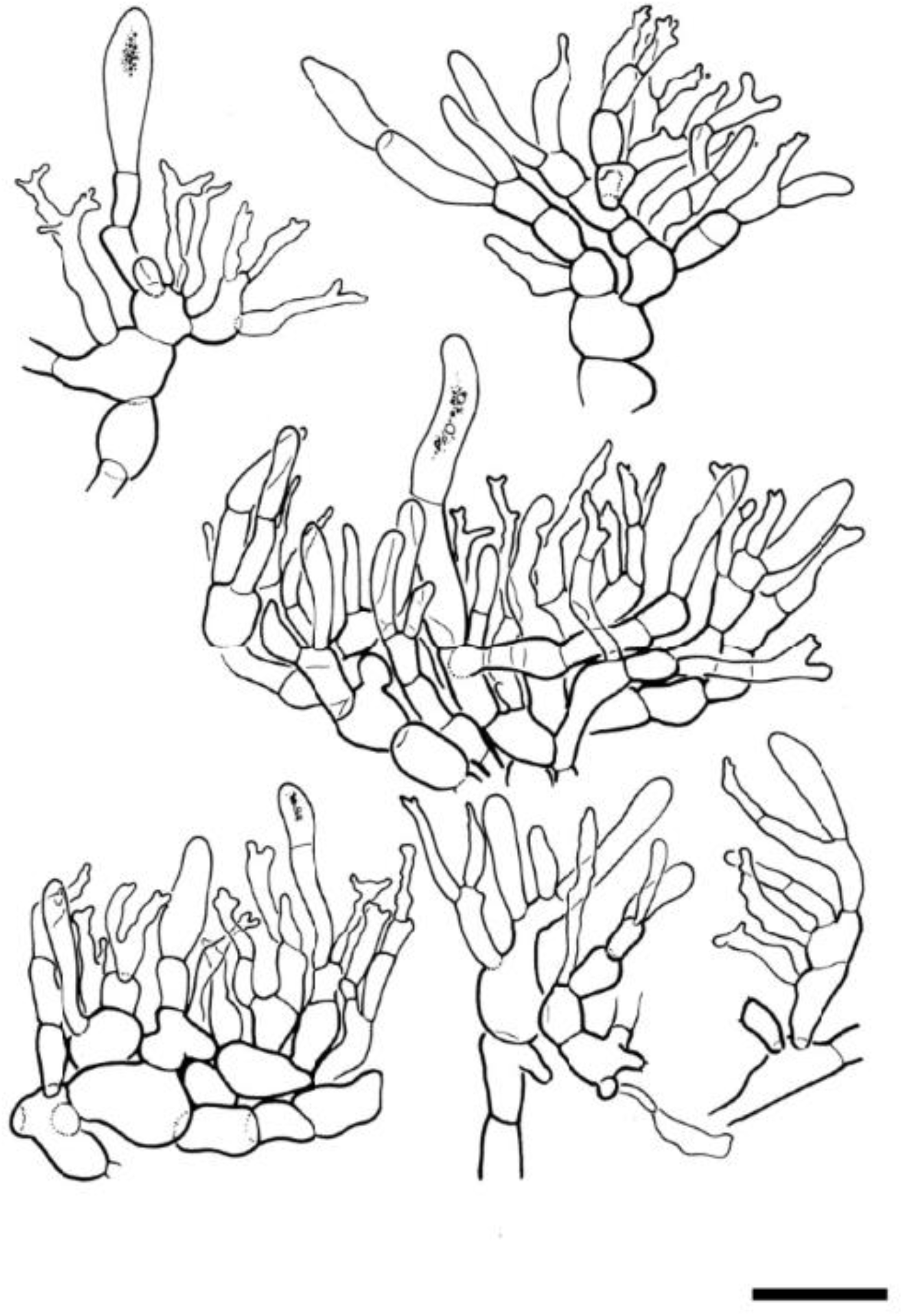
*Russula koniamboensis* Buyck, sp. nov. (*722/BB09.022*, holotype). Fragments of the suprapellis. - Scale bar = 20 µm. Drawings B. Duhem.

**Fig. 14.**
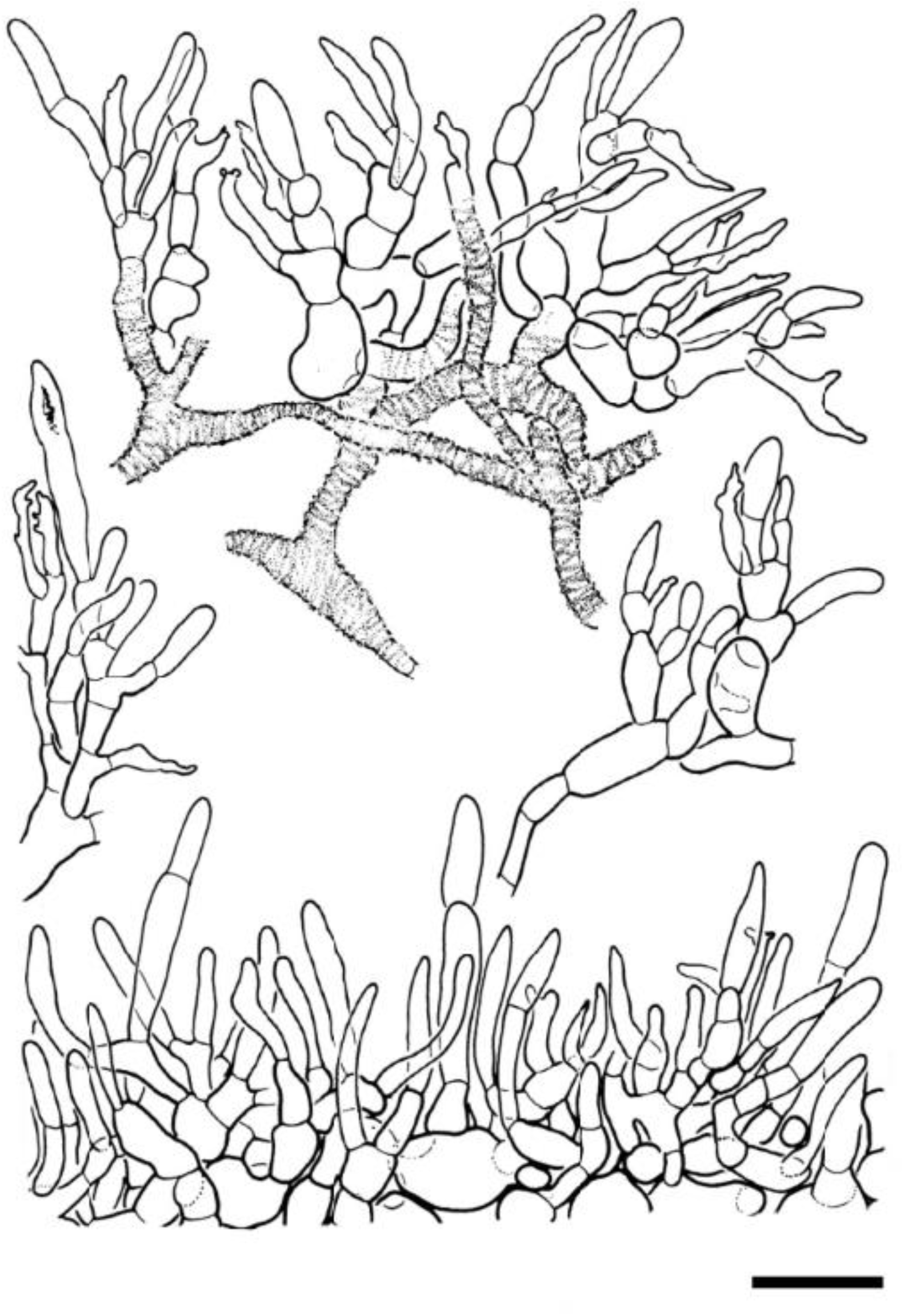
*Russula koniamboensis* Buyck, sp. nov. (*730/BB09.117*): Details of the suprapellis. - Scale bar = 20 µm. Drawings B. Duhem. *Notes*: The above description is based on the holotype collection. However, the two other collections of this species (having identical ITS sequences) clearly show that its microscopic features are quite variable among different collections. For the pileipellis this variation concerns the size of the sphaerocytes in the lower suprapellis which are sometimes up to 35 µm diam., giving rise to 8-celled chains (as in *BB09.346*); observed variations apply also to the presence/absence of distinct orthochromatic zebroid incrustations on the cell walls in the lower pileipellis (present in *BB09.117*) and finally also to the form of the terminal cells on the hyphae. The latter can be either more regular in outline and similar to those of *R. incrustata* (as in *BB09.346* and *BB09.117*) or strongly diverticulate as in the type collection. On the lamella edge, cheilocystidia can be abundant and well-developed (as in *BB09.346*) or just dispersed among the other elements. The spores in *BB09.346* are near-identical in size and ornamentation to the holotype [(8.13)8.20-**8.63**-9.05(9.38) x (6.25)6.47-**6.86**-7.26(7.50) μm, Q = (1.11)1.17-**1.26**-1.35(1.40)], while spores in *BB09.117* produced several aberrant sizes and ornamentations indicating an unfavourable development. These spores were, therefore, not used for measurements.

Differs from *R. incrustata* in the pale yellowish pileus, white stipe, the somewhat larger spore size and its occurrence under *Nothofagus*; from *R. purpureotincta* principally in the pileus colour and its geographic distribution.

*Holotype*. — New Caledonia. Northern Prov., Massif du Koniambo, near Voh, in the nickel mine exploitation site called ‘Niko’, S 21* 00’22’’ – E 164*49’51’’, at 724 m alt., on ultramafic soil under *Nothofagus balansae*, 17 March 2009, leg. *B. Buyck, 722/BB 09.022* (PC0142407).

Mycobank. —

*Additional examined material*. — New Caledonia. Northern Prov., Massif du Koniambo, near Voh, 734 m alt., in the “île nickel” mine exploitation site, S 21* 00’42’’ – E 164*49’50’’, on ultramafic soil under *Nothofagus balansae*, 19 March 2009, leg. *B. Buyck 730/BB 09.117* (PC0714855); Massif du Koniambo, near the Trazy entry of the nickel mine exploitation site, on ultramafic soil under *Nothofagus codonandra*, 9 April 2009, leg. *B. Buyck, 742/BB 09.346* (PC0714856).

Etymology. — named after the type locality.

#### Pileus

25-30 mm diam., convex, slightly depressed in the center, near the margin striate over 1/3-1/4 of the radius, surface dull, smooth to finely or even distinctly farinaceous in the center and more or less concentrically deposited, peeling 2/3 of the radius, pale yellowish in the center, cream towards margin.

#### Lamellae

Equal in length, adnate, distant and ca. 1-2 L/mm at pileus margin, off-white to cream color, obtusely rounded at the pileus margin; entire edge concolorous.

#### Stipe

30-34 x 6-8 mm, central, cylindrical to slightly obclavate, glabrous, smooth to finely longitudinally striate, white to ivory, pale greyish towards base, brittle, spongy inside, lacunes absent.

#### Context

Very thin toward the margin, white, distinctly greying in age. Taste and odor not distinctive.

#### Spore print

white.

#### Spores

(6.67)7.51**–8.01**–8.51(8.75) x (6.25)6.52–**6.83**–7.15(7.29) μm, Q = (1.06)1.10–**1.17**– 1.24(1.31), subglobose to broadly ellipsoid; ornamentation subreticulate, composed of large, prominent, moderately distant, conical to hemispherical and strongly amyloid spines, up to 1(–1.5) μm long, connected by dispersed to frequent fine lines into a (very) incomplete network; suprahilar spot well–developed, varying from strongly amyloid to verruculose and grayish to poorly amyloid.

#### Basidia

30–42 × 12–17 μm, clavate, of variable shape, fusiformous to distinctly clavate, with (2–)4 stout sterigmata; basidiola clavate.

#### Subhymenium

pseudoparenchymatic.

#### Hymenial gloeocystidia

68–94 × 7–12 μm on lamellar sides, smaller near the lamella edges, up to 45 μm long, narrowly clavate to fusiform or subcylindrical, frequently mucronate to appendiculate at the apex, up to 14 μm long, originating in subhymenium and protruding beyond the basidia, thin– walled with walls up to 1 µm thick; contents in Congo Red mainly restricted to dispersed refringent inclusions of variable size that do not react to sulfovanillin.

#### Marginal cells

15–26(–34) × 4–6(–9) μm, sitting on 1–2 short, subterminal cells, small, occupying most of the lamellar edges, extremely variable in shape, similar to the smaller terminal cells in the suprapellis in having 1–4 diverticulate, obtuse lobes or outgrowths.

#### Pileipellis

40–60 μm thick, two–layered, not well delimited from the underlying context, in the lower part composed of hyphae with or without distinct orthochromatic incrustations, forming a pseudoparenchyma of intertwined and strongly ramifying, ascending to erect, irregularly shaped, thin–walled cells; basal cells sometimes up to 20(–25) µm wide, near the surface giving rise to short chains composed of 2–4(–5), narrower cells that are slightly inflated and barrel–shaped, ellipsoid to subcylindrical, up to 10 µm wide; terminal cells of variable size and either longer or shorter than subterminal cells, (8–)16–23(–34) × 2–4(–6) μm, often very irregular or undulate in outline, slightly attenuating toward the frequently capitate, appendiculate diverticula at apex. Primordial hyphae recognizable mostly by their somewhat more regular outline, but especially by the refringent granular–heteromorphic contents of the terminal cell that mostly measures 15–25 × 4–5 μm, narrowly clavate to subcylindrical in outline, obtuse rounded at the apex, diverticula or appendages absent, thin–walled. Cystidioid hyphae absent in subpellis and context. Oleiferous hyphae rare.

#### Clamp connections

absent.

### 3. Russula purpureotincta R.F.R. McNabb, New Zealand Journal of Botany 11: 711. 1973

Figs. 15-18

**Fig. 15.**
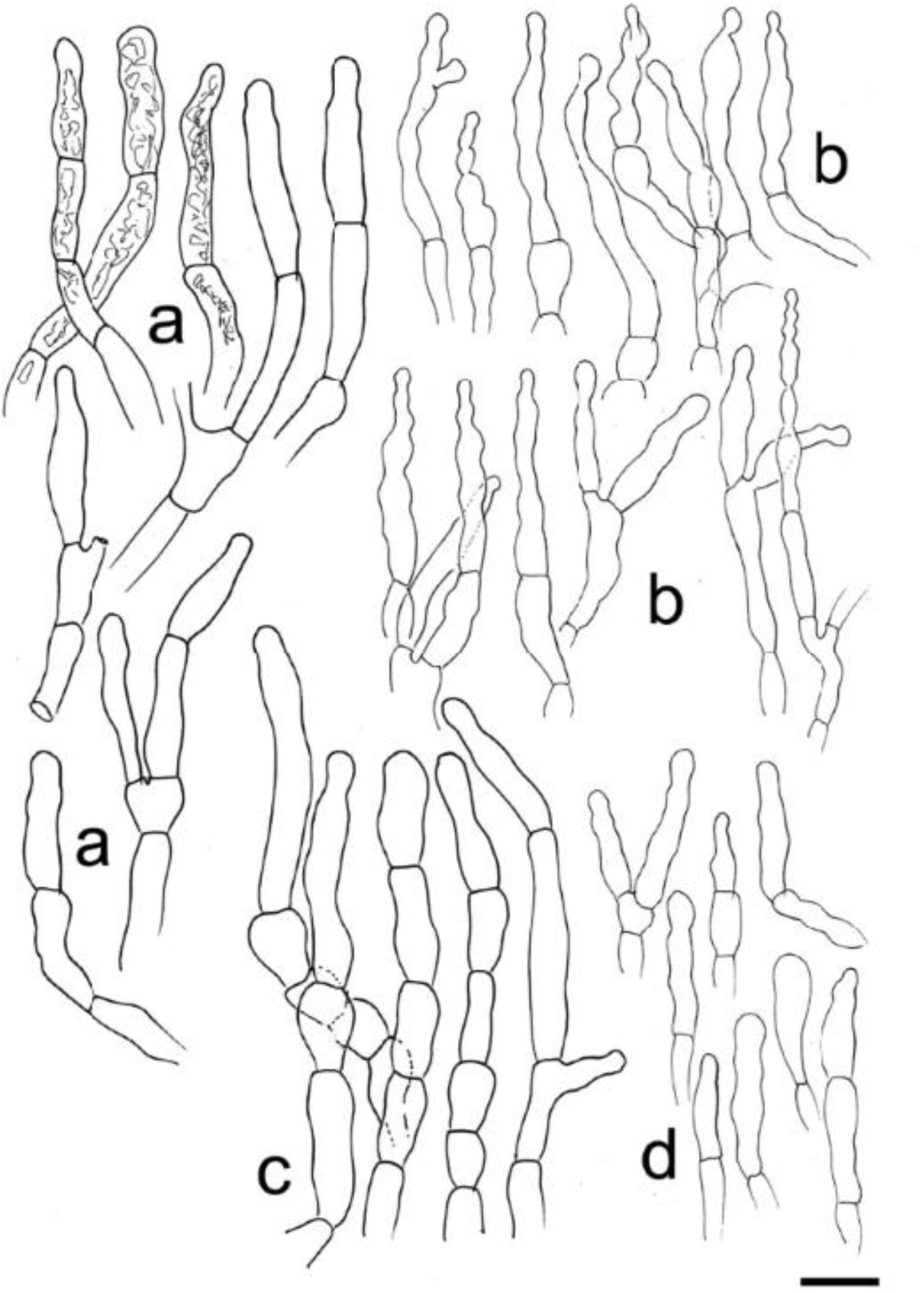
*Russula purpureotincta* (*ZT 68-105*): a-b. Elements of the pileipellis near pileus margin: a, primordial hyphae with indication of contents in few cells; b, terminal cells of the other hyphal extremities; c-d. Elements of the pileipellis in the pileus center: c, primordial hyphae without indication of contents; d, terminal cells of other hyphal extremities. Scale bar = 10 µm (Drawings B. Buyck)

**Fig. 16.**
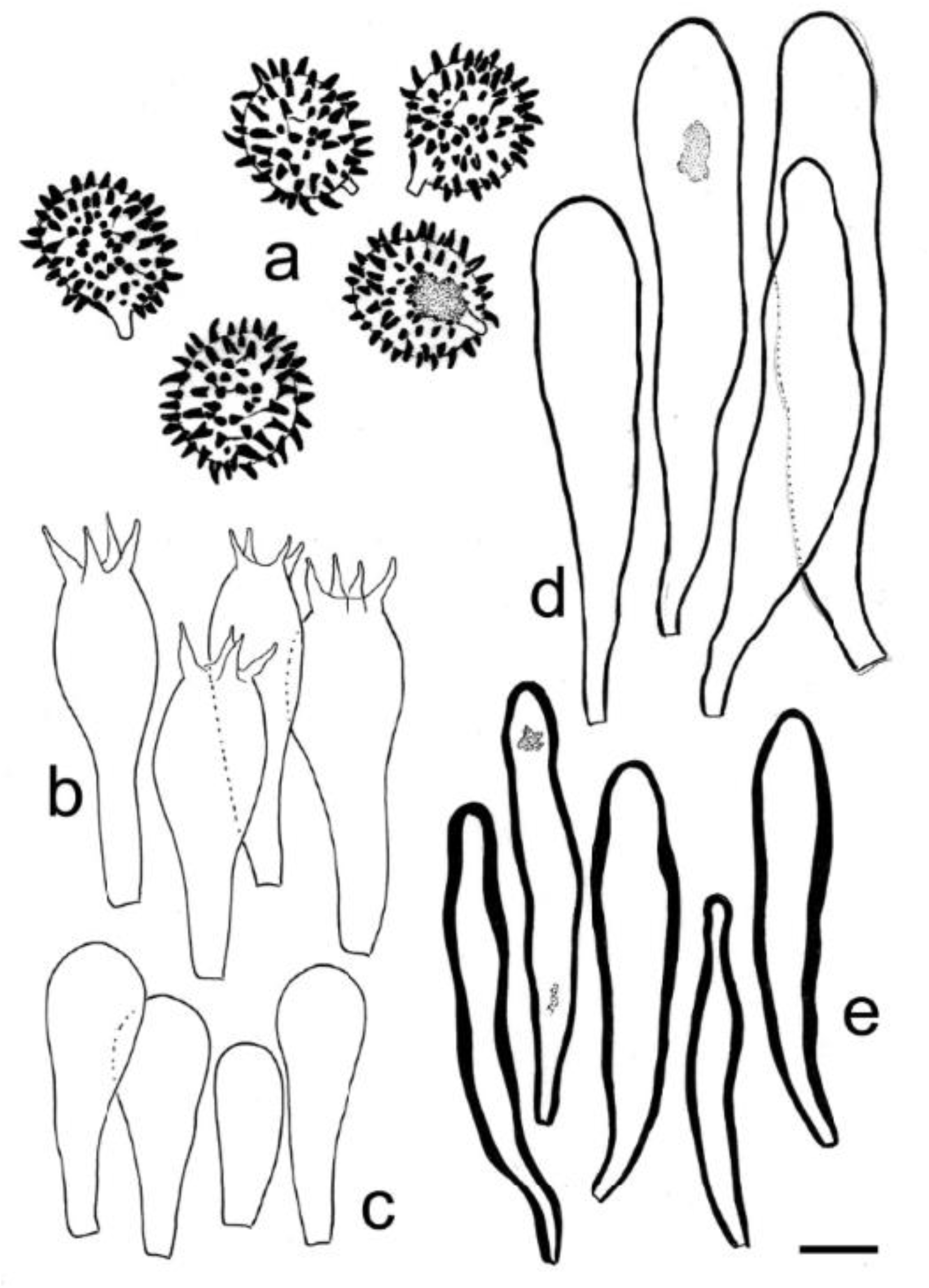
*Russula purpureotincta.* Elements of the hymenium (all from *ZT 68-105*, except b from *ZT 68-086*). a-b, basidiospores; c, basidia; d, basidiola; e, pleurocystidia; f, cheilocystidia. Scale bar = 10 µm. Drawings B. Buyck.s.

**Fig. 17.**
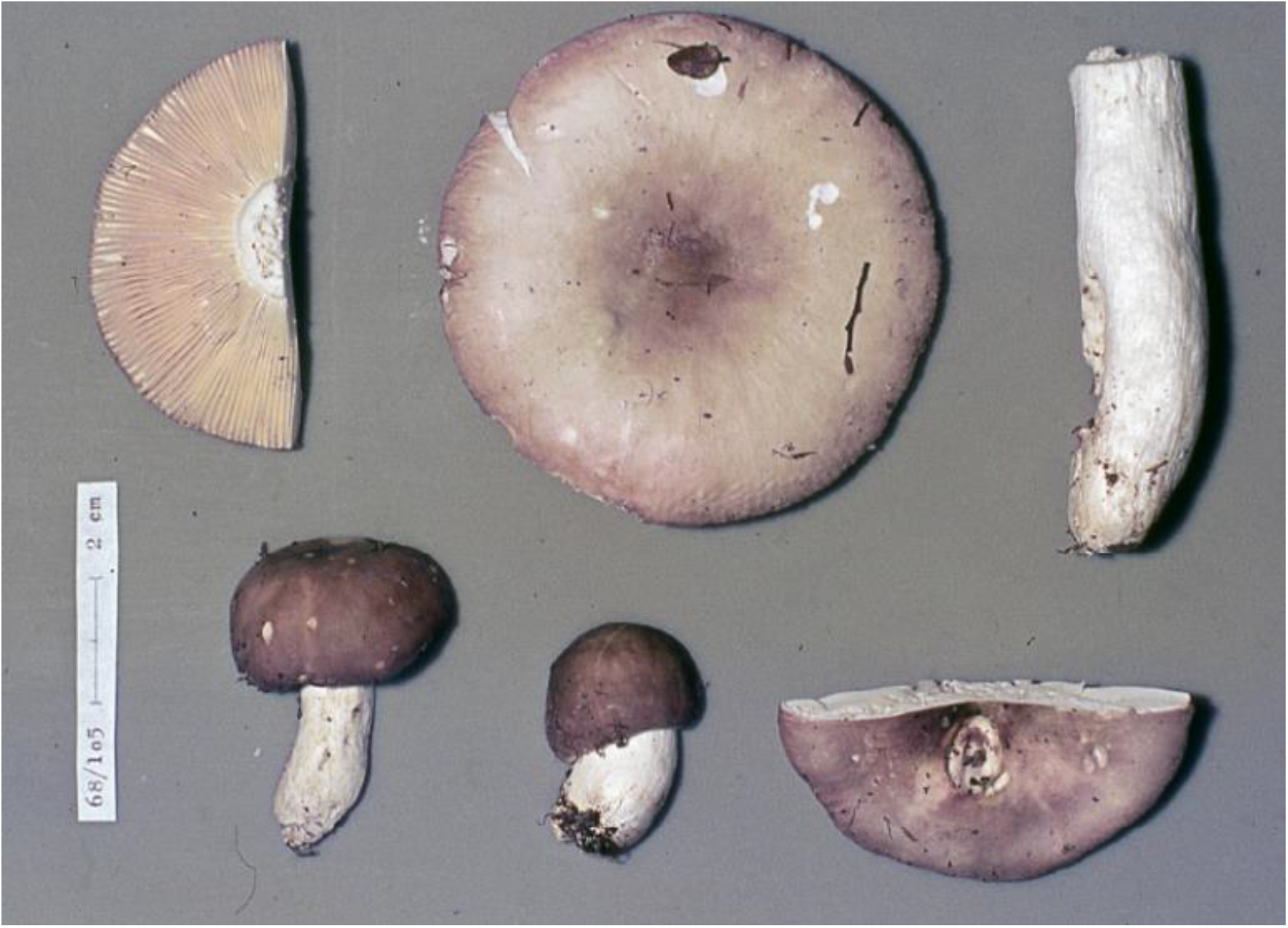
*Russula purpureotincta*. Fresh basidiomata (*ZT 68-105*). Photo E. Horak.

**Fig. 18.**
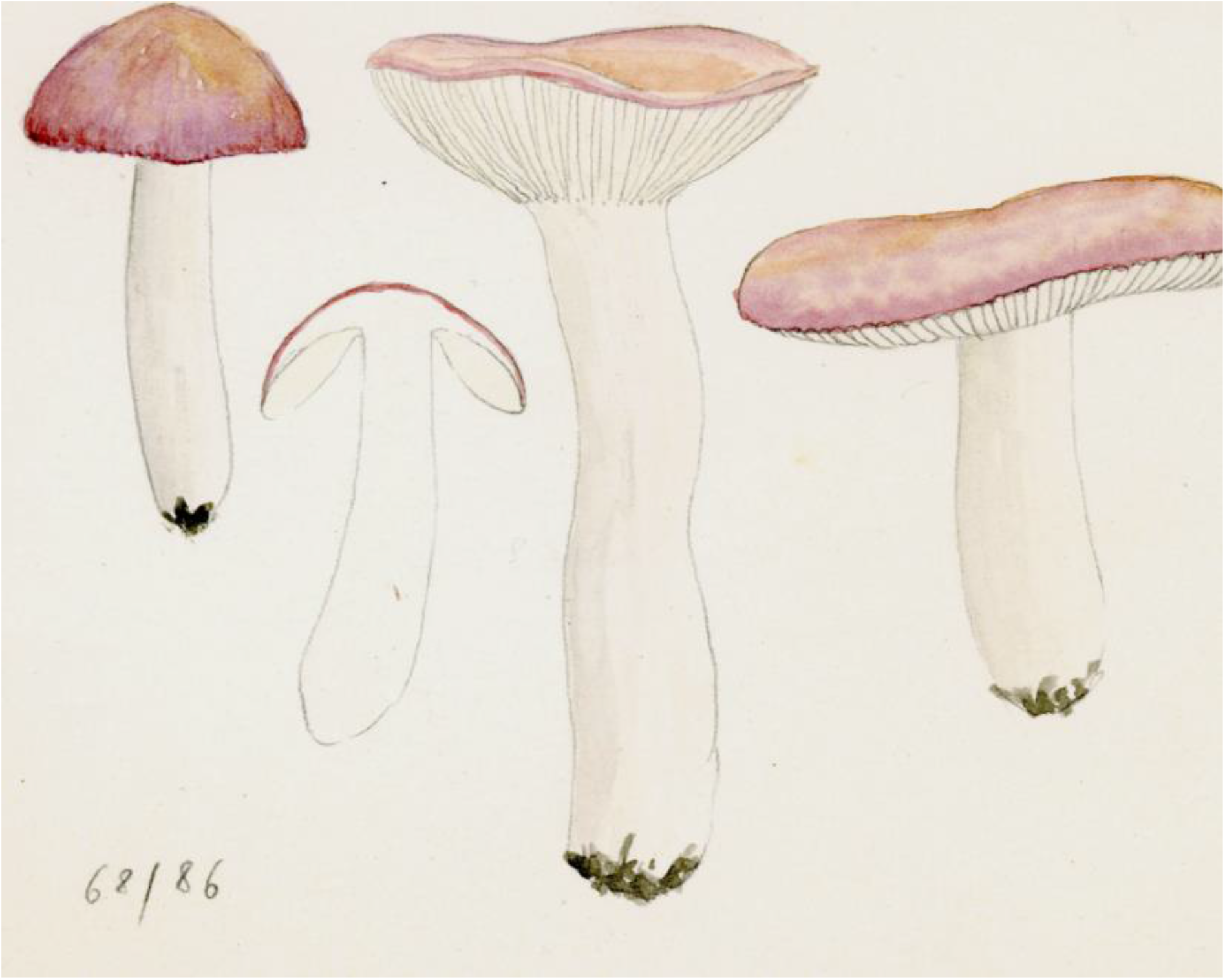
*Russula purpureotincta*. Fresh basidiomata (*ZT 68-068*). Watercolour by E. Horak.

*Examined material*. — **New Zealand**. Prov. Nelson, Springs Junction, Upper Grey, among *Sphagnum* under *Nothofagus [Fuscospora] solandri* var. *cliffortioides* and *Nothofagus [Lophozonia] menziesii*, 25 Febr. 1968, *E. Horak (ZT 68-086)*; Prov. Nelson, W of Tophonse Saddle, at the border of a swampy locality between moss (not *Sphagnum*) under *Nothofagus [Fuscospora] solandri var. cliffortioides* and *Nothofagus [Lophozonia] menziesii*, 3 March 1968, *E. Horak (ZT 68-105)*.

#### Pileus

Up to 75 mm diam., at first convex, becoming slightly depressed in the center with age, near the margin smooth to shortly striate; surface dull, when humid weakly viscid, at first reddish lilac to purplish, but rapidly discoloring, particularly between the center and the very margin, and then becoming beige-brownish-pinkish to flesh-colored, pileipellis separable up to mid-radius.

#### Stipe

36-80 x 10-19 mm, cylindrical or slightly narrowing towards apex, entirely white, smooth or finely striate lengthwise.

#### Lamellae

Equal in length or with rare lamellulae, 4-6 mm wide, normally spaced (ca. 1L/mm at the pileus margin), white, becoming cream with age, not anastomosing in the dorsal interspaces, obtusely rounded at the pileus margin; gill edge smooth, concolorous.

#### Context

brittle, spongy, white, unchanging on exposure. Odour not distinctive. Taste mild.

#### Spore print

white.

#### Spores

(8.13–)8.32–***8.74–8.93***–9.43(–10.21) x (6.67–)6.99–***7.28–7.29***–7.57(–7.92) µm, Q = (1.11–)1.14–***1.20–1.23***–1.28(–1.34), large, subglobose to broadly ellipsoid, densely ornamented with strongly to partly amyloid spines or cylindrical warts, these up to 2 µm high in many spores and frequently curved, often in pairs, with few smaller granular warts and rare interconnecting fine lines; suprahilar spot distinctly amyloid.

#### Basidia

40–48 x 12–15 µm, clavate, 4–spored; sterigmata stout, mostly 7–8 x 2–3 µm.

#### Hymenial gloeocystidia

Mostly 70–90 x 12–15 µm, abundant, on lamellar sides clavate to fusiform, thin– to slightly thick–walled; near lamellar edges distinctly thick–walled (up to 2–3 µm thick in middle portion) and smaller, mostly 40–60 x 7–12 µm, rarely with an additional septum in the upper part.

#### Marginal cells

Not differentiated.

#### Pileipellis

Two–layered: the suprapellis forming a trichoderm of densely packed, narrow and thin– walled hyphal extremities composed of 3–6 short cells and originating from a pseudoparenchymatic layer underneath; the latter is composed of much more inflated, sausage–like, ellipsoid or spherical cells up to 30 µm diam., without zebroid incrustations on the cell walls; suprapellis composed of hyphal extremities with terminal cells in the pileus center short to very short, rarely exceeding 20 x 3–5 µm, either narrowing in the upper part or clavate, often slightly undulate in outline, toward the pileus margin more irregular in outline and very frequently globose to moniliform at apex. Typical pileocystidia absent. Primordial hyphae emerging from the surface of the suprapellis, slightly more voluminous than the other hyphal extremities and composed of 3–6, thin-walled cells, usually more strongly septate in the pileus center; their cells shortly cylindrical to barrel–shaped, and mostly also recognized by distinctly refringent, granular contents, up to 8 µm diam., the terminal cell frequently up to 35 µm long, more regular in outline compared to the other sometimes subcapitate hyphal extremities. Cystidioid and oleiferous hyphae lacking.

#### Clamp connections

absent.

*Notes*: The description is based on two collections gathered by E. Horak one year before the species was described by R.F.R. McNabb (1973) as *R. purpureotincta*. Our description is highly similar to the original description, taking into account that the spore size given by McNabb (9-12 x 8.5-10.5 µm) includes the spore ornamentation. In addition, McNabb was probably not aware of the concept of ‘primordial hyphae’.

This endemic species of New Zealand seems to be widely distributed and is not rare as many records are reported online (e.g. on https://scd.landcareresearch.co.nz). The online available field images, illustrations of microscopic features and SEM pictures of the spores for these collections (including for the type collection) clearly confirm our own observations on Horak’s specimens.

*Russula purpureotincta* is represented in our ITS phylogeny by nine sequences, including one newly produced sequence from the type specimen and one [GU222259 for PDD77740] that corresponds to a very pale collection initially identified as *Russula cremeo-ochracea* (and still deposited as such in GenBank). McNabb’s species resembles our *R. incrustata* because of the faint greyish magenta, greyish purple to reddish grey tints present in the pileus, but it lacks the warmer orange-red tints of our New Caledonian species which, in addition, has a coloured stipe. As evident from online published pictures for *R. purpureotincta*, the stipe can frequently be obclavate and can vary considerably in length between different specimens.

*Russula purpureotincta* is very variable in overall colour, but typically discolours very rapidly leaving a pale sordid whitish-isabelline pileus with only faint greenish-pinkish hues remaining in the center and at the very margin of the pileus. Such discoloured specimens are more reminiscent of equally discoloured forms of several (but acrid !) species in the *Russula* core clade, rather than of other species of subsect. *Roseinae*.

Microscopic differences between *Russula purpureotincta* and both above-mentioned New Caledonian species are very subtle. Spore ornamentation in McNabb’s species seems less reticulate, its cheilocystidia more strongly thick-walled, and the contents in hymenial cystidia more abundant.

## Discussion

Subsect. *Roseinae* is sister to subsect. *Lilaceinae* in all recent multigene phylogenies. Both subsections have always been considered – even in the traditional morphotaxonomic classifications (Romagnesi, 1967; Sarnari, 2003) – either as one single or as two extremely close species groups. In older morphology-based classifications, both subsections were part of subgen. *Incrustatula* Romagnesi (1967) together with other subsections having more coloured spore prints. In the first multigene phylogeny that was more or less representative of world-wide *Russula* species, Buyck et al. (2018) demonstrated the artificial concept of subgen. *Incrustatula* with dark-spored subsect. *Amethystinae* (Romagn.) Bon and *Chamaeleontinae* Singer of *Russula* subgen. *Incrustatula* being unrelated to the pale-spored taxa in subsect. *Roseinae* and *Lilaceinae*. This was later confirmed by the same authors in Rossi et al. (2020) when providing significant, although moderate support to group the northern hemisphere, dark-spored subsections *Amethystinae* and *Chamaeleontinae*, together with the Oceanian subsection *Tricholomopsidum* Buyck & V. Hofst. and the Asian subsect. *Castanopsidum* Buyck & X. H. Wang and not with the *Roseinae* – *Lilaceinae* clade.

Secotioid or hypogeous *Roseinae* have not yet been reported from the northern hemisphere. We here place two secotioid southern species in the subsection: *R. [Macowanites] kermesina* from New Zealand and *R. albobrunnea* T. Lebel from Australia. Our ITS phylogeny suggests that these southern secotioid taxa are related to different lineages of northern hemisphere *Roseinae* compared to the agaricoid Roseinae from New Caledonia and New Zealand (Fig. 1).

The southern *Roseinae* associate rarely with host trees from family Myrtaceae (so far only *R. incrustata*) and seem to prefer *Nothofagus* as a host (*R.purpureotincta, R. kermesina, R. koniamboensis, R. albobrunnea*). It is not clear yet whether these hosts have been more recently invaded by these fungi or not but, so far, no *Roseinae* have been reported from *Nothofagus* forests in South America.

Arguing that the traditional interpretation of *Roseinae* should be considered at the section level, Looney et al. (2022) described subsect. *Albidinae* Looney, Manz & Adamčik, for merely two species (*Russula albida* and one still undescribed species). The same authors also suggested that a separate subsection was needed for the American /magnarosea clade, another tiny subclade based so far on two undescribed collections. In our opinion, proposals to elevate such extremely small entities within terminal clades to higher ranks should be postponed until more worldwide samplings for the various *Russula* clades become available.

Buyck et al. (2018) noticed that many of the southern hemisphere *Russula* have a pseudoparenchymatic pileipellis structure. This observation was based on the fact that all of species in the hyperdiverse subsect. *Tricholomopsidum* have a pseudoparenchyma in the lower suprapellis. The latter feature also applies to all *Roseinae* that are discussed here. It is yet unclear whether such a pileipellis structure represents an advantage for the involved species.

## Acknowledgements

B. Buyck obtained funds for field work in New Caledonia both by ANR (French National Agency for Research), BIONEOCAL ANR-07-BDIV-006 (PI P. Grandcolas, CNRS-MNHN) and ULTRABIO ANR-07-BDIV-010 (PI M. Ducousso, CIRAD). Sequencing of the collections was financed through a 2014 project submitted to the ATM « Emergences » (Dir. S. Peigné) of the National Natural History Museum in Paris. The first author is also thankful to the late B. Duhem for the microscopic drawings of the New Caledonian species. Mr. Denis Poignonec of Koniambo Nickel SAS is thanked for access to the Koniambo Massif mining area. Mr. David Paulaud of the “Direction de l’Environnement” for Southern Prov. is thanked for the collecting permits in New Caledonia. E. Horak is grateful for logistic support offered by the New Zealand Forest Research Institute (Rangiora) and the Herbarium PDD in Auckland. Song acknowledges funding from the Research Initiation Project of Shaanxi University of Technology (SLGRCQD2214) and the Science and Technology Department of Shaanxi Province project (2022JQ-199).

